# Epitope Effect Prevalence in Affinity-based pQTL studies

**DOI:** 10.1101/2025.06.20.660695

**Authors:** Jurgis Kuliesius, Mihaly Badonyi, Pau Navarro, Joseph A. Marsh, Lucija Klaric, James F. Wilson

## Abstract

Affinity-based proteomics platforms Olink and SomaScan have enabled population-wide human proteogenomic studies, linking genetic variants to protein abundances. However, sequence alterations in the binding site of the detection reagent may introduce platform-specific measurement bias unrelated to protein levels, referred to as the epitope effect. In this study, we investigated the prevalence of epitope effects using *cis* protein quantitative trait loci (pQTL) discovered in three of the largest proteogenomic studies: UK Biobank, deCODE, and Fenland.

Across 5,817 protein targets assayed in these studies, *cis*-pQTL were identified for 914 proteins by both platforms, 301 (33%) of which were linked to a missense variant. We identified 37 proteins with opposing effect directions in two platforms for the same missense pQTL, and 85 proteins where a missense pQTL was significant in only one platform.

We present examples where such discrepancies reflect differences in isoform or proteoform targeting, as well as examples where the discordance appears to result from true platform-specific detection bias. Further structural analyses reveal that missense *cis*-pQTL are more likely to be detected when they alter residues located on accessible protein surfaces - regions most likely to interfere with reagent binding in affinity-based assays.

Overall, our findings suggest that missense-mediated epitope effects influence only a minority (12% or less) of *cis*-pQTL results. We also highlight the need for detailed assay annotations and structural context to improve result interpretation in proteogenomic studies.

## Introduction

Protein quantitative trait loci (pQTL) analyses, where pQTL are identified through genome-wide association studies (GWAS), have been instrumental in linking genetic variation to circulating protein levels, providing insights into disease mechanisms 1, drug targets2, and biomarker discovery^3^. Recent advances in affinity-based proteomics, particularly the Olink^4^ and SomaScan^5^ platforms, have enabled large-scale pQTL studies, collectively profiling thousands of proteins across diverse populations^6, 7, 8^. However, platform-specific discrepancies and technical artifacts pose challenges in interpreting these associations and their biological relevance9, 10. These include differences in pQTL detection, effect size, or effect size directionality for the same protein across platforms, complicating the interpretation of whether observed associations reflect true biological variation. Mass spectrometry-based approaches are also vulnerable to biases arising from protein-altering variants, as differences in peptide sequence and their associated mass-to-charge ratios can lead to incorrect protein quantification^1^.

Missense variants at the affinity-reagent binding site, or epitope, have the potential to alter the binding of the aptamer (SomaScan) or antibody (Olink). Previous proteogenomic studies have thus speculated that such epitope effects are a potential source of bias in results obtained from affinity-based proteomics technologies^10, 11, 12, 13^. In such cases, the protein measurement would reflect increased or decreased binding of the aptamer or antibody, rather than the intended abundance of the protein. The resulting epitope effect pQTL may be indistinguishable from genuine pQTL when using standard proteogenomic approaches^14^.

Given the potential of epitope effects to influence pQTL signals, they offer a valuable opportunity to assess both biologically meaningful regulatory effects and assay-specific biases. While these concerns are supported by mechanistic reasoning, empirical evidence quantifying the extent of such effects across platforms has been limited to disease trait colocalization^2^, pQTL comparisons to eQTL^3^, and matching affinity pQTL to mass-spectrometry assays^4^, which have their own biases. We provide an extension of previous methods by integrating structural epitope features to investigate their potential to interfere with reagent binding. To do this, we analysed *cis* missense pQTL identified through GWAS using two high-throughput affinity-based proteomic platforms, comparing and contrasting results for the proteins quantified with both3, 4, ^12^.

In this study, we analysed sentinel *cis* pQTL data from three of the largest proteomics studies by sample size (UK Biobank, deCODE & Fenland) to investigate missense variant-driven pQTL associations. By harmonizing datasets across platforms and studies, we identified that up to 36% of proteins with a *cis*-pQTL have at least one missense pQTL in high LD with a lead variant within the gene encoding the targeted protein. We then examined platform-specific inconsistencies in missense pQTL detection and effect size directionality, hypothesizing that such discrepancies could arise from epitope effects rather than true biological regulation. To further characterize the structural impact of these variants, we incorporated predictions of Gibbs free energy change (ΔΔG, a measure of the change in protein stability), and surface accessibility derived from AlphaFold^15^ protein structures, to assess potential interference with antibody and aptamer binding.

Through this comprehensive approach, we aim to refine the interpretation of pQTL signals, disentangle true regulatory effects from assay-related biases, and provide a framework for improving the reliability of affinity-based proteomics interpretations in research.

## Results

### Missense *cis*-pQTL prevalence

The three largest European proteomic GWAS to date - deCODE and Fenland (analysed using SomaScan) and UK Biobank (Olink) - collectively assay a proteomic breadth of 5,817 human protein targets in plasma, with up to 4,720 via the SomaScan v4.0 platform and 2,923 on the Olink Explore panel [Supplementary Table 1]. Of these, 1,827 proteins (39% of those measured by SomaScan) and 1,861 proteins (64% of those measured by Olink) have been reported to have at least one *cis-*pQTL in these studies, thereby suggesting a genomic confirmation of successful capture of the target molecule^10^. Among these, 914 protein targets have at least one genome-wide significant *cis*-pQTL in both proteomics platforms [Fig. 1].

**Figure 1.**
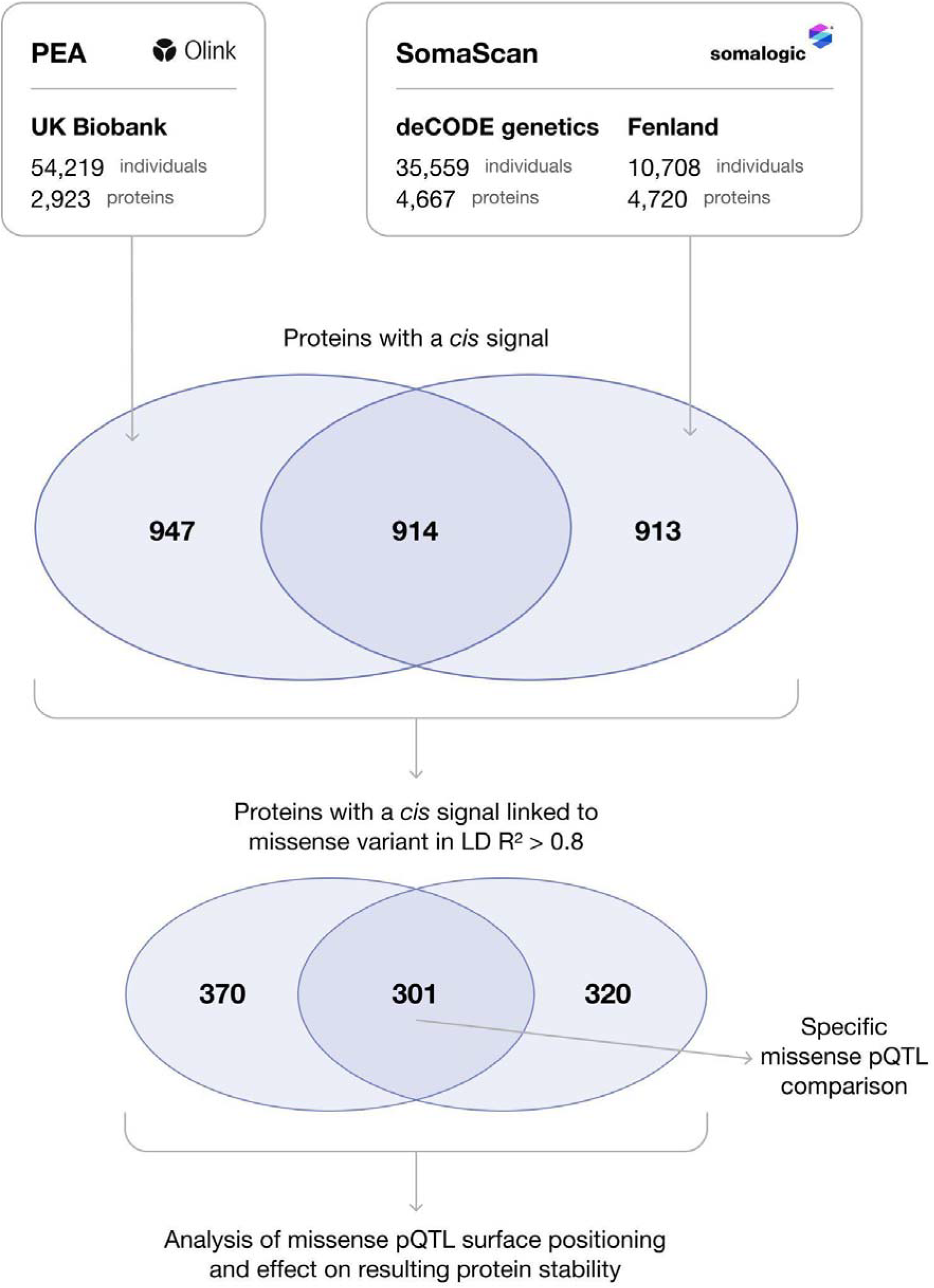
Diagram of the filtering steps used to identify proteins with *cis* pQTL linked to missense variants and subsequent analyses. This study used publicly available data from the UK Biobank (Olink), deCODE genetics, and the Fenland (SomaScan) studies.

To assess which of the *cis-*pQTL are in LD with a missense variant and therefore potentially artefacts caused by epitope effects, we annotated each independent *cis* pQTL to include all proxy variants within LD (R^2^ > 0.8). We then filtered this expanded set of correlated variants at each independent pQTL for missense variants. Among proteins with *cis*-pQTL, 621 (34%) of proteins in SomaScan and 671 (36%) in Olink have one or more missense variants in strong LD with at least one of the reported independent *cis* variants [Supplementary Table 2].

### Contrasting measurements

Among proteins with *cis* pQTL detected on both proteomic platforms, 301 (33%) exhibited at least one independent missense variant-associated pQTL in either or both assays. For 190 proteins with *cis* missense pQTL, no obvious discrepancies across platforms were apparent in terms of direction or evidence of a pQTL effect. On the other hand, among 111 proteins, we identified two distinct patterns of disagreement between the two platforms, with some proteins showing both. Firstly, 37 proteins showed genome-wide significant missense *cis*-pQTL in both platforms, but with opposing effect directionality [Supplementary Table 3, Supplementary Figure 1]. Secondly, 85 proteins, each with at least one *cis*-pQTL in both proteomics platforms, but with one or more independent significant missense *cis*-pQTL in studies using one platform but failing to reach genome-wide significance for those missense *cis-*pQTL in studies using the other proteomics platform [Supplementary Table 4]. To illustrate the nature of platform-specific disagreement in missense *cis*-pQTL effect sizes, we examined individual cases where effect direction or detection differed between the two proteomic technologies. In all cases, allele frequencies were closely matched between studies (R2 > 0.98), in accordance with their shared Northwest European heritage [Supplementary Fig 5].

Among the 37 proteins with opposing effect directions for a particular genome-wide significant *cis* missense variant, one of the most striking examples was rs1859788 (chr7:99971834 A>G), a missense variant in the *PILRA* gene. This variant showed both exceptionally strong statistical associations (p < 3.7x10^-3376^ in UK Biobank and up to 1.2×10^-27511^ in SomaScan studies) and some of the largest effect size magnitudes within this discordant pQTL subset [Supplementary Figure 1]. rs1859788 encodes a change from arginine (positively charged, bulky side chain) to glycine (smallest, neutral amino acid) in the 78^th^ position and has been previously shown to considerably affect the binding kinetics of both endogenous and pathogen-derived ligands16. Located in the second exon, the variant is present in all three splice variants of *PILRA* that are the named targets of three distinct aptamers in the SomaScan v4.0 (5k) assay – the canonical sequence, and two isoforms FDF03-deltaTM and FDF03-M14, with missing or altered transmembrane domains, respectively [Supplementary Table 1]. Notably, while rs1859788 was identified as the leading missense variant for decreasing the circulating levels of both truncated isoforms (FDF03-deltaTM deCODE β = 1.31, p < 1.2×10^-27511^, Fenland β = 1.26, p <1.1×10^-5297^; FDF03-M14 deCODE not analysed, Fenland β = 1.27, p < 5.7x10-5244), it had no association with plasma levels of the membrane-bound form of PILRA in the SomaScan assay (deCODE p < 0.70; Fenland p < 0.75) [Supplementary Figure 2]. In contrast, the UK Biobank study using the Olink assay found rs1859788 to be strongly associated with increased circulating levels of PILRA (β = -1.00, p < 3.7×10^-3376^).

Another similar example can be found with Plasminogen and rs143079629 (chr6:161128812 G>A), a missense variant at *PLG* causing replacement of arginine with lysine in the 89th position of the encoded protein. SomaScan measures three distinct proteoforms of PLG with three separate aptamers – plasminogen, plasmin and angiostatin – all of which encompass the 89^th^ amino-acid within their structures. Plasminogen is the precursor of plasmin while angiostatin is generated by the proteolytic cleavage of either plasminogen or plasmin by proteases, including plasmin itself17. Of these three proteoforms, only circulating angiostatin was associated with rs143079629 in SomaScan. Plasminogen measurements were associated with a different missense *cis*-pQTL, rs4252129, that was not in LD with the discordant rs143079629 (LD R^2^ = 0.0002) in both SomaScan studies, while the measurement of plasmin had no genome-wide significant associations in the *cis* region of *PLG*. Meanwhile, the UK Biobank Olink study reports 4 independent *cis* signals for their Plasminogen measurement, with the same missense variant rs143079629 being genome-wide significant and in high LD with one of them (rs537579467, R2 = 0.96) [Supplementary Figure 3]. rs143079629 was found to substantially decrease the circulating levels of angiostatin when analysed using SomaScan but it increased the levels of plasmin in the study using the Olink platform without having a measurable effect in SomaScan.

Of the 85 proteins where a genome-wide significant missense *cis*-pQTL was captured by only one of the two proteomics platforms [Supplementary Table 4], we highlight results for two protein measurements for which there are no known alternate proteoforms annotated in the UniProt database^18^ (accessed May 2025). This selection was made to ensure that the observed platform-specific discrepancies are not driven by differences in proteoform targeting, allowing us to focus on other potential sources such as epitope effects or assay-specific biases. Complement factor H-related protein 5 has been found to have increased circulating levels with an arginine-to-histidine substitution at position 356, mediated by the genomic variant rs35662416 in the *CFHR5 gene* (chr1:196967354 G>A). This effect was exclusively observed in the two studies using the SomaScan platform (p = 1.6×10^-818^ and p = 1.3×10^-133^), but this signal was entirely absent in Olink (p = 0.45). We note that additional independent *cis* signals with no correlated missense variants were also observed in both SomaScan and Olink data for this protein, suggesting that while the missense pQTL is specific to the SomaScan platform, the protein itself is robustly detected by both assays [Figure 2].

**Figure 2.**
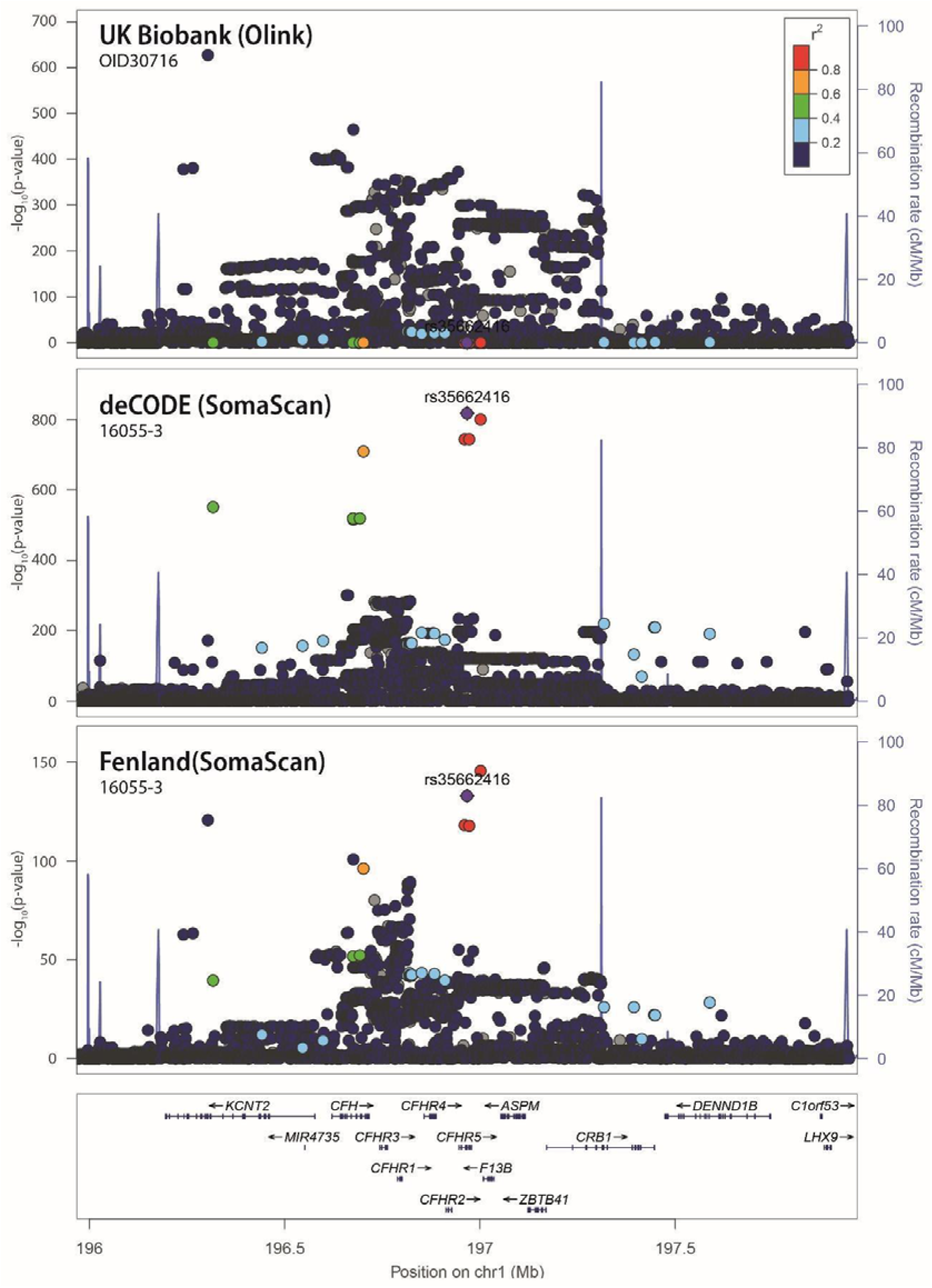
LocusZoom plots of missense rs35662416 associations with *CFHR5* measurements across three proteomic studies. rs35662416, a missense pQTL for *CFHR5*, shows strong associations in both SomaScan studies (deCODE and Fenland), but the signal is entirely absent in the UK Biobank study using the Olink platform. This discrepancy suggests a platform-specific detection bias, potentially due to an epitope effect. The LD pattern in each LocusZoom plot is colour-coded with respect to the missense pQTL.

Missense variant-driven platform discrepancy was also observed for Serine protease inhibitor Kazal-type 6 (SPINK6), a small, 80-amino-acid protein primarily involved in regulating skin barrier function through the inhibition of epidermal proteases19. rs12186491 (chr5: 147593497 C>A) was identified as either the lead SNP or in strong LD (R^2^ = 0.98) with one in both studies using the SomaScan technology (p = 2.6×10^-1235^ and p = 3.4×10^-537^). Meanwhile, the similarly powered UK Biobank study also reported a strong association in *cis* with *SPINK6* using the Olink assay (rs12717962, p = 1.5×10^-531^), a pQTL also independently genome-wide significant in the SomaScan studies (rs12186491 - rs12717962 LD R^2^ = 0.06). The association between circulating levels of *SPINK6* and rs12186491, however, was not supported by the Olink assay (p = 0.02) [Supplementary Figure 4].

### Missense pQTL surface positions and effect on protein stability

Next, we sought to determine whether the location of the altered residue (buried within the tertiary protein structure or surface-exposed) contributes to the likelihood of detection as a missense *cis*-pQTL. This is of relevance for affinity-based platforms, where changes in epitope accessibility may play a crucial role in detection molecule binding affinity and impact measurement sensitivity. To this end, we analysed the structural context of missense *cis*-pQTL by integrating data from AlphaFold-predicted protein structures. We used relative solvent-accessible surface area (RSA) as a quantitative measure to assess how exposed each replaced residue is within the folded protein structure. To facilitate interpretation, we grouped residues into three RSA-based structural classes: buried (RSA < 0.2), representing residues with minimal solvent exposure typically located in the protein core or hydrophobic regions; intermediate exposure (RSA 0.2 – 0.65), corresponding to residues situated on the protein surface; and highly exposed (RSA > 0.65), often found in unstructured or disordered regions of the protein^20, 21^.

Investigating all missense *cis-*pQTL reported in the Olink and SomaScan platforms, we observed a greater proportion of amino acid substitutions occurring on the protein surface, when compared to the genome-wide average distribution for missense variants identifiable in the referenced proteomics studies, given their power to detect such variants (MAF > 0.001) [Figure 3, Supplementary Table 5]. Fisher’s exact test showed that missense pQTL in both Olink (odds ratio = 1.32, p-value = 3.1×10^-3^) and SomaScan (odds ratio = 1.41, p-value = 4.2×10^-4^) have a similar moderate enrichment in the intermediate 0.2 – 0.65 RSA range, compared to the baseline. This suggests that missense *cis*-pQTL are more likely to be detected when they affect surface-exposed residues, as opposed to buried residues or those in disordered regions, across both Olink and SomaScan proteomics platforms.

**Figure 3.**
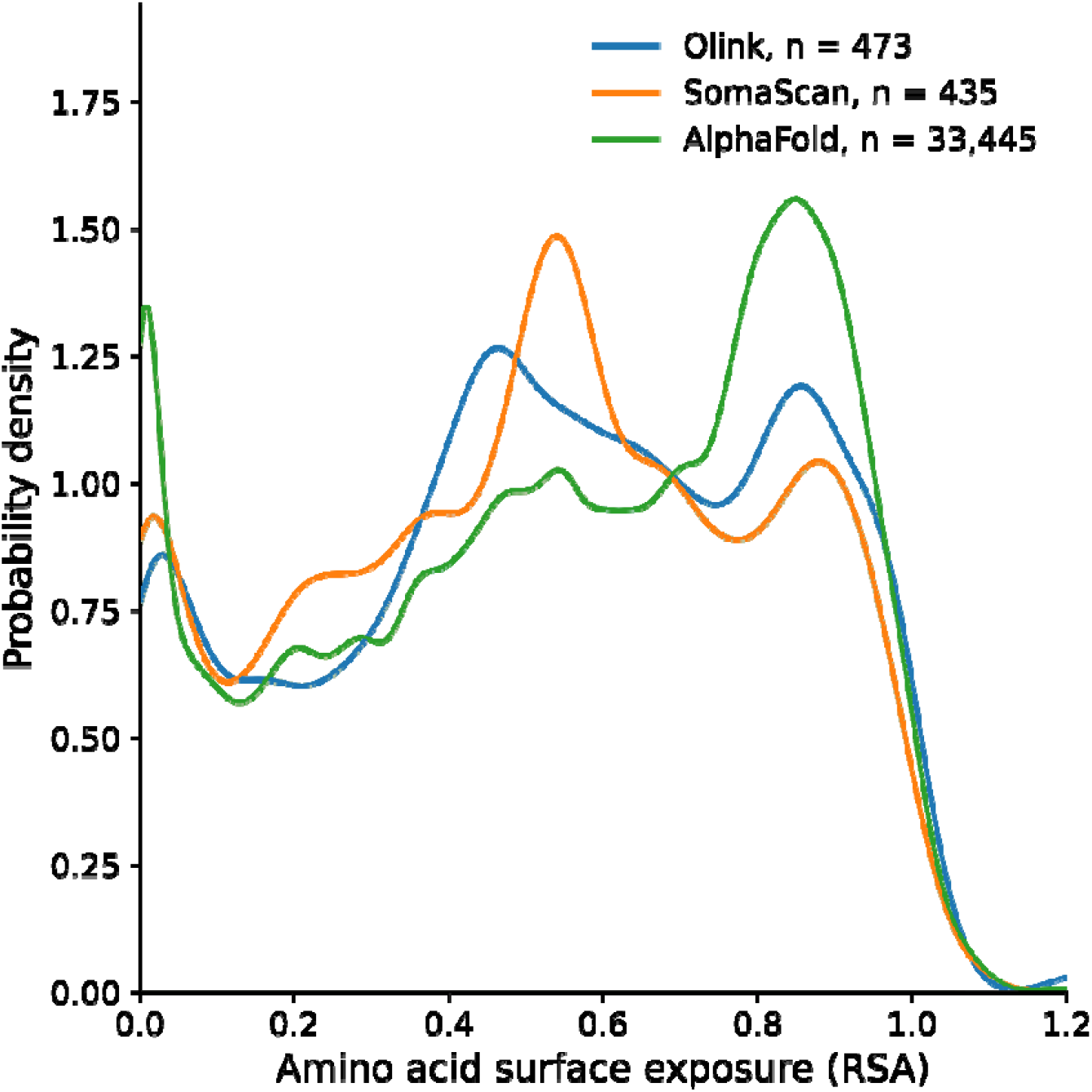
Distribution of relative solvent accessibility (RSA) for missense variants across datasets. The density plots reveal that missense *cis*-pQTL in both Olink and SomaScan proteomics platforms are modestly enriched in moderately exposed surface regions (RSA 0.2– 0.65). This suggests that missense variants influencing protein levels via *cis*-pQTL mechanisms are more likely to occur in structured, surface-accessible regions. This is possibly due to the greater likelihood of interfering with detection reagent binding by modifying the epitope in affinity-based assays. Data by colour: Olink *cis* missense pQTL (Blue); SomaScan *cis* missense pQTL (Orange); All detectable (MAF > 0.001, Global) missense variants (Green).

To explore systemic mechanisms contributing to platform-specific detection discrepancies, we tested whether missense variants affecting surface charge were disproportionately represented among discordant *cis-*pQTL. This was motivated by the possibility that the negatively charged DNA aptamers used in SomaScan may be sensitive to changes in target surface charge, as previously suggested^13^. Among the 39 missense *cis-*pQTL with platform-discordant associations in effect direction [Supplementary Table 3], 19 were located on surface-accessible regions of the protein (RSA 0.2–0.65). Of these, 8 involved a substitution that resulted in the loss of a positively charged residue (arginine, lysine, or histidine). This was significantly enriched compared to the background of surface-exposed missense variants from proteins measured by both platforms (odds ratio = 3.54, p = 9.2×10^-3^). In contrast, genetic variants leading to the gain of positive charge were not enriched among discordant variants (p = 0.65). Repeating the analysis for platform-discordant pQTL significant in only one platform but not the other [Supplementary Table 4], we found no evidence of enrichment for either the loss (p = 0.071) or gain (p = 0.40) of positive charge among surface-exposed residues. This shows that while loss of positive surface charge may underlie some of the observed platform-specific detection differences, it does not appear to be a widespread mechanism underlying all platform-specific differences.

To further explore the structural consequences of missense *cis*-pQTL, we assessed their impact on protein stability by calculating the missense-mediated protein structure changes in Gibbs free energy (ΔΔG), relative to the corresponding wild-type proteins. To this end, we have performed a ΔΔG rank concordance analysis between missense *cis* pQTL detected using Olink and SomaScan platforms and the detectable missense variant subset [Figure 4]. If structural deformation were a key mechanism driving epitope interference, we would expect an overrepresentation of high ΔΔG ranks in our pQTL subset. However, no link between the identified pQTL and their effect on the protein’s structural stability was observed in the Olink and SomaScan platforms, suggesting that missense variant-linked structural protein change magnitude does not systematically affect pQTL detection.

**Figure 4.**
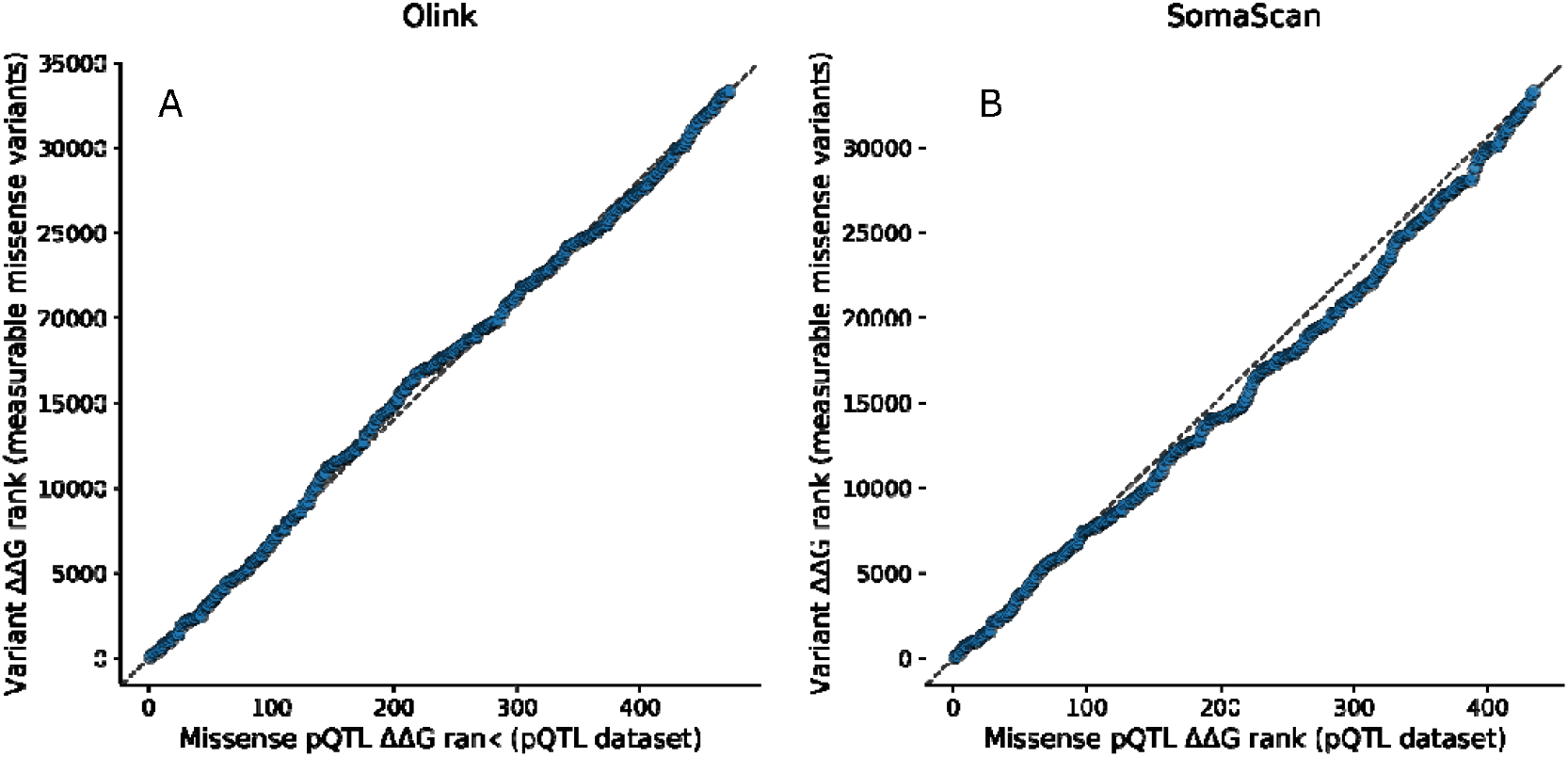
pQTL effects on protein structural change have no connection to the observed platform-specific differences. The scatter plots show no deviation from the mean in the ranked predicted ΔΔG values (change in Gibbs free energy upon mutation) of missense *cis* pQTL variants in Olink (A) and SomaScan (B) proteomic platforms. x-axis: Rank of ΔΔG for *cis* missense pQTL in Olink (A) or SomaScan (B); y-axis: Rank of ΔΔG for the same missense variants between all detectable (MAF > 0.001, Global) missense variants. Diagonal dashed line represents perfect agreement in rank between datasets.

## Discussion

In this study, we utilise the presence of a *cis-*pQTL in a proteomic GWAS as strong evidence that the proteomic assay is measuring the intended protein, rather than a spurious molecule (even if there could still be a technical artifact)^10^. This is because genetic variants located near or within the gene encoding a given protein have a far higher likelihood of influencing that protein’s abundance than variants elsewhere in the genome. Consistent with previous research10, we have confirmed the successful targeting of a similar number of proteins in genetic studies for both Olink and SomaScan assays (1,861 vs 1,827), even though the proportion of proteins with one or more *cis* signals is much greater for Olink (64% vs 39%).

By taking advantage of the Northwest European descent of the three studies, we were able to draw on the LD structure of a subset of genetically British UK Biobank individuals to expand our search for missense variants beyond those reported as lead independent pQTL in the original studies. Our analysis revealed that a substantial proportion of proteins with *cis*-pQTL - 36% (671 proteins) in Olink and 34% (621 proteins) in SomaScan – were associated at a genome-wide level with at least one *cis* missense variant, either directly as a lead pQTL (626 pQTL, 51%) or through proxy variants in strong linkage disequilibrium (LD R^2^ > 0.8) (595 pQTL, 49%) [Supplementary Table 2]. This finding suggests that measurements of up to a third of proteins with a *cis* signal may be vulnerable to biases directly caused by epitope-altering genetic variants.

Among the 914 protein measurements with *cis*-pQTL detected across both SomaScan and Olink platforms, we identified a substantial subset (301 proteins, 33%) linked to at least one independent lead *cis* missense variant, enabling a direct comparison of missense pQTL between the two technologies. While the majority of these signals (63%) were concordant across platforms, the remaining discrepancies may reflect potential epitope effects. Two distinct patterns of disagreements between the platforms emerged: opposing effect directionality for the same missense *cis*-pQTL (observed for 37 proteins) and platform-specific significance (85 proteins with missense *cis*-pQTL reaching genome-wide significance in only one of the two platforms). Allele frequencies of the platform-specific missense were consistent between the studies [Supplementary Figure 5].

Some of the observed discrepancies between the SomaScan and Olink platforms can be explained by the two targeting different protein isoforms. In one instance, PLG is measured with three separate aptamers in the SomaScan platform, with each aptamer measuring a different proteoform – plasminogen, plasmin and angiostatin – each with unique biological functions and properties^17^. One of these measurements shared a missense *cis*-pQTL with Olink’s measurement of PLG, but with opposite effect sizes. A similar scenario was observed for PILRA, where SomaScan also had three distinct aptamers targeting three isoforms of the protein. Olink had only one measurement of the protein that shared a missense *cis*-pQTL with one of SomaScan’s measurements of PILRA. Closer inspection revealed that in both cases the SomaScan assay was capturing variant proteoforms, while the Olink assay likely reflected the canonical form of the protein. This difference in assay annotation provides a plausible explanation for some of the observed pQTL discordance.

In contrast, we share two other case studies of discordant results for CFHR5 and SPINK6, proteins with no known alternate isoforms or proteoforms, which showcase true platform-specific pQTL detection. In both instances, missense *cis*-pQTL were robustly detected by one platform but were absent in the other, despite comparable statistical study power. Our results imply three key points: (1) even in the absence of proteoform complexity, epitope effects can lead to platform-specific detection artefacts; (2) genome-wide significant *cis-*pQTL signals may reflect changes in binding affinity rather than true protein abundance; and (3) the absence of pQTL signal in one platform does not guarantee biological irrelevance. While we cannot with certainty claim that all 85 proteins with platform-specific missense *cis-*pQTL are affected by epitope effects, such discrepancies suggest that both antibody- and aptamer-based assays can exhibit distinct binding affinities influenced by missense variants and highlight the importance of orthogonal validation when interpreting platform-specific pQTL.

Given that relatively little is known about the epitope sites that aptamers bind to^22^, we then investigated whether amino acid changes caused by missense pQTL detected using the aptamer-based SomaScan platform and the antibody-based Olink platform differ from the proteome-wide distribution of missense variants. Since antibodies are known to bind structured, surface-exposed regions of proteins23, 24, 25, we hypothesized that Olink-associated missense pQTL would be overrepresented in such residues. If highly destabilizing missense variants were indeed contributing to epitope effects, we would also expect to observe a skew toward larger than average changes in protein stability among missense pQTL. Annotating each missense pQTL with the relative solvent-accessible surface area (RSA) from the resulting amino acid substitution, we find that variants discovered through both proteomics platforms are similarly enriched in residues that are solvent-exposed and reside in structured protein domains. Furthermore, we found that missense variants causing a loss of arginine, lysine, or histidine on the outer protein surface were significantly enriched among effect-direction discordant pQTL, suggesting that disruption of electrostatic interactions may contribute to aptamer^13^ or antibody^25^ binding changes in a subset of cases. Our findings suggest that despite their differing chemistries, both aptamers and antibodies appear to be similarly prone to epitope effects due to amino acid change on accessible, structured surfaces.

Significant alterations in protein structure caused by missense genetic variants were also considered as potential drivers of epitope effects. Using the predicted change of Gibbs free energy (ΔΔG), we assessed the impact of the missense mutations at (or in strong LD with) *cis*-pQTL on the resulting protein stability by calculating ΔΔG values relative to their canonical counterparts. No enrichment in highly stabilizing or destabilizing amino acid changes was observed in the pQTL identified through either Olink or SomaScan technologies. Thus, missense pQTL-mediated conformational shifts, as estimated by ΔΔG, are unlikely to be an important mechanism driving epitope effects and observed association differences between Olink and SomaScan.

Taken together, our results demonstrate that although what are likely to be epitope effects contribute to the observed discrepancies between protein measurements obtained through SomaScan and Olink platforms, the overall impact of changes in epitope caused by missense variants is modest, affecting at most 12% of currently assayed proteins. In particular, only a subset of proteins (37%) with genome-wide significant pQTL linked to a missense variant for the measured protein exhibited platform-specific associations or opposing effect directions attributable to possible epitope interference. For the majority of matched protein targets with a *cis* signal (88%), either no missense variant was associated with the measured protein, or the missense variants had consistent effects that were detected by both technologies. While some concordant missense associations could still be a result of an epitope effect — if both the aptamer and antibody bind to the same site on the protein surface — this likely accounts for a minority of such associations. This suggests that biological variation in protein abundance, not measurement artifacts caused by epitope effects, is the main driver of the observed *cis*-pQTL signals. While this study emphasizes the importance of considering epitope effects in pQTL interpretation, it also provides reassurance that both SomaScan and Olink affinity-based proteomics assays predominantly capture genuine genetic influences on protein levels.

In the absence of identifiable epitope sites, proteomics technology providers should offer detailed target annotations for each assay. This should include the specific protein isoform targeted, the amino-acid sequence of the protein that the antibody or aptamer was selected against, and any post-translational modifications. Such information would greatly improve the interpretation of platform-specific associations and the identification of potential causes of detection bias.

Meanwhile, proteogenomics researchers can assess whether a given missense *cis* pQTL obtained through either SomaScan or Olink is at risk of causing an epitope effect by referencing the affected amino acid residue in the protein’s AlphaFold-predicted structure. This study has shown that missense variants located in structured, surface-accessible regions with loss of positive charge are more likely to interfere with reagent binding than those buried within the protein core or situated in intrinsically disordered regions. Structural context can thus be a valuable tool for distinguishing biological signals from assay-specific artifacts.

All structural, enrichment and contrast analyses in this study were restricted to proteins with at least one *cis-*pQTL in both SomaScan and Olink platforms. While this approach enabled direct cross-platform comparisons, it excluded proteins uniquely measured by one platform, those lacking a *cis* signal in one of the assays, and those not measured by either platform. In all these cases epitope effects may also be prevalent but remain unanalysed. As a result, our conclusions about the structural properties of missense *cis*-pQTL may not generalise to the full set of platform-specific targets, and the overall burden of epitope effects could be either under- or overestimated.

In this study, we used AlphaFold2-predicted structures, which models proteins in their monomeric forms. However, many human proteins function as part of larger complexes^26^, and residues that appear surface-exposed in monomeric predictions may become occluded when the protein is assembled into its physiological complex. We have not assessed the prevalence of missense pQTL within protein-protein interfaces, but we emphasize the need for precise target annotation in affinity-based proteomics. Specifically, clarity on whether aptamers or antibodies bind the monomeric form of the protein, the protein as part of a physiological complex, or all circulating forms indiscriminately without bias would greatly improve the interpretation of proteogenomic results. Such information should ideally be provided by assay manufacturers or inferred through bioinformatics approaches like those applied in this study.

Another limitation of our study is that even with both affinity-based platforms showing concordant results for a particular missense variant, they may both be affected and binding to the same epitope. This raises the possibility that some of the concordant associations may still reflect assay artifacts rather than true change in protein abundance associated with the genotype. As such, orthogonal validation using mass spectrometry is essential to confirm whether these associations represent genuine biological variation or are driven by assay-specific biases.

A further limitation of our study is that we did not account for other genetic variants that can directly impact protein structure, such as stop-gained or frameshift mutations. Additionally, we did not consider post-translational modifications, like glycosylation, methylation, and proteolytic cleavage, which can significantly alter protein surfaces and epitope accessibility^27^. These modifications can thereby lead to variability in protein detection even in the absence of missense genetic variation. As a result, some observed pQTL signals — or lack thereof — may reflect changes in protein modification states other than the *cis* missense variants investigated here.

Finally, our study assumes that the missense *cis* variant in highest LD with the lead variant is the most likely candidate driving each observed association signal. However, this may not always be the case, as the lead pQTL could be in LD with another variant affecting gene expression, splicing, or a regulatory element. Without fine-mapping or functional validation, it remains possible that the association is due to another variant in the region, and attributing the signal solely to the missense change may oversimplify the underlying genetic architecture.

Despite these limitations, our study highlights the value of integrating proteogenomic and structural protein data at scale, enabling more nuanced interpretations of *cis-*pQTL signals across technologies. By systematically identifying missense pQTL-linked platform discrepancies, this work contributes to improved quality control and interpretation of proteogenomic results. Our approach demonstrates that structural protein features can be feasibly incorporated into large-scale pQTL analyses without requiring bespoke experimental data, adding an orthogonal layer of insight to aid prioritising candidate variants for functional follow-up.

Looking ahead, we offer a practical framework for identifying and interpreting potential epitope-driven artifacts in proteogenomic studies. Our results indicate that most platform-concordant signals likely reflect genuine biological variation rather than measurement artifacts, and that both technologies are similarly sensitive to surface-exposed epitope-altering variants. As affinity-based proteomics continues to expand in coverage of the human proteome, increasing awareness of epitope effects will improve the robustness of protein–trait associations and accelerate the translation of proteogenomic discoveries into clinical and therapeutic applications.

## Methods

### pQTL collection

We analysed publicly available protein quantitative trait loci (pQTL) data from three major affinity-based proteomics studies: (1) Olink measurements of 2,923 proteins in 54,219 UK Biobank participants^4^, (2) SomaScan analysis of 4,667 proteins in 35,559 Icelandic individuals from the deCODE genetics study^3^, and (3) SomaScan measurements of 4,720 proteins in 10,708 participants from the Fenland study^12^. We selected these studies as they focused on individuals of the same (Northwestern European) ancestry and offered comparable statistical power of detecting robust genetic associations. To ensure comparability across the three studies and minimize annotation mismatch, we standardized GWAS summary statistics across the three datasets. Protein measurements between different proteomics platforms were matched using UniProt identifiers and standardized to HGNC gene symbols. Genomic variants were remapped to GRCh37 coordinates using USCS’s LiftOver tool^28^ where necessary, while pQTL effect size direction and allele frequency data was reoriented to a common effect allele. To ensure unambiguous mapping of *cis* genetic regions and to maintain consistency across the different studies and proteomic platforms, we excluded detection molecules annotated to bind more than one protein target. An overview of the analysed proteins, including whether they were measured across the different assays, as well as a summary of identified *cis* pQTL and missense variant-associated *cis* pQTL for each protein, is provided in the Supplementary Table 1.

### Missense *cis* pQTL identification

We defined missense associations as variants that: (1) were annotated as missense in gnomAD v2.129, and (2) were missense for the gene encoding the protein measured by the proteomics assay. For each of the three studies, we downloaded complete summary statistics for all protein measurements that showed significant *cis* pQTL associations in the respective study.

We extracted variants within ±1 Mb of each target protein’s transcription start site for the ensuing LD-based clumping analysis. To ensure consistency across studies with different imputation panels, we harmonized datasets by retaining only SNPs that: (1) had rsID annotations, and (2) were present in all three studies. This filtering step had a mean of 54.7% of retained SNPs across all protein measurements (range: 2.3-92.0%; SD: 23.3%; see Supplementary Table 6).

For exceptionally strong GWAS associations (-log10(p) ≥ 300), Fenland and deCODE studies reported the p-value as p = 0 or -log10(p) = infinite. In these instances, we have obtained the chi-squared statistic (*χ*^2^) from the estimated variant effect size and its standard deviation, and then derived the p-value using the programmatic expression below:

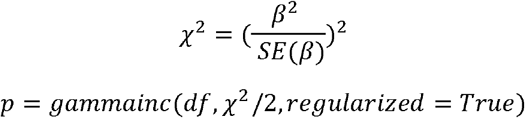

*χ*^2^ - Chi-squared statistic, estimating whether β is significantly different from 0. Note that Z^2^ = *χ*^2^ with one degree of freedom.

β - The estimated effect size from the GWAS regression model. SE(β) - The standard error of β.

p – p-value, representing the probability under the null hypothesis of observing a *χ*2 statistic as extreme or more extreme than the computed value.

gammainc: The regularized incomplete gamma function, which computes the cumulative probability for a *χ*^2^ distribution. Function implementation used from the mpmath 1.3.0 module in python 3.9.18.

df - Degrees of freedom (df=1 for single-SNP tests in GWAS).

### LD-based missense pQTL prioritisation

Given that all studies were conducted in European-ancestry populations, we performed linkage disequilibrium (LD)-based clumping using LD patterns from a 10,000-individual White British subset of UK Biobank as the reference panel^30^ using PLINK 1.90^31^. We applied a Bonferroni-corrected genome-wide significance threshold for the GWAS p-value (p = 5x10-8), further corrected for the 2893 autosomally encoded proteins with a reported *cis* association that were analysed in this study (p = 1.73×10^-11^). We have used a stringent LD correlation metric (R^2^ = 0.001) to ensure picking up only fully independent signals within 1 Mb up and downstream of the transcription start site of the encoded protein. Specifically, clumping parameters used were --clump-p1 1.73e-11; --clump-r2 1e-4; --clump-kb 1000.

To link the *cis* pQTL signals to potential missense-mediated epitope effects, we next searched for missense variants in high LD (R^2^ > 0.8) with each lead variant. For each protein’s independent *cis* pQTL, we subsequently identified any missense variants in high LD using PLINK 1.90 using the same 10,000 White British subset of UK Biobank as the reference panel. PLINK parameters used were --r2; --ld-window-r2 0.8; --ld-window-kb 1000 --ld-window 9999. If one or more missense variants were found to be in high LD (R^2^ > 0.8) with a lead pQTL, we prioritized the missense pQTL showing the strongest LD with the lead variant as the most plausible candidate for assessing missense-mediated epitope effects (column Missense SNP, Supplementary Table 2). Conversely, pQTL lacking any known *cis* missense variants in high LD were considered unlikely to be driven by an epitope effect and were therefore excluded from further analysis.

### Platform-specific missense discrepancies

The following criteria were used to identify missense *cis* pQTL as potential indicators of epitope effects, assay-specific biases, or other technical artifacts between the SomaScan and Olink assays:

The two assays having identified the same lead missense *cis* pQTL as genome-wide significant but with opposing effect size directionality;

Both assays detected at least one *cis* pQTL for a given protein, but a lead missense *cis* pQTL was genome-wide significant (p < 1.73×10^-11^) in one assay while non-significant in the other.

Protein measurement assays fulfilling any one of these criteria were labelled as discordant and as potentially affected by an epitope effect. These missense variants were then selected for a separate, more detailed investigation [Supplementary Table 2].

### Relative solvent accessibility

Relative solvent-accessible surface area (RSA) quantifies the extent to which an amino acid residue is exposed to the solvent within the context of the folded protein. It is defined as the ratio between the solvent-accessible surface area of a residue within the monomeric protein and the maximum solvent accessibility observed in a Gly-X-Gly tripeptide, where X represents the amino acid of interest32. This normalization allows for comparison across different residues by accounting for their inherent structural constraints.

RSA values were calculated with AREAIMOL from the CCP4 programme^33^ suite using AlphaFold34-predicted protein structures15. These were used as reference in order to quantify and compare solvent exposure distributions. Of the >5.5 million missense variants present in gnomAD v2.1^29^, 33,445 variants with global minor allele frequencies (MAF) higher than 0.001 as identified in either 1000 Genomes^35^ or GnomAD Exomes^29^ (dbSNP build 156^36^). These variants were located within 5,831 encoded proteins measured as single-protein targets in studies using either SomaScan or Olink assays. The MAF threshold was selected as a conservative estimate of the lower limit of variant detection given the sample sizes of included studies, with the lowest reported pQTL MAF being 0.0001 in the deCODE SomaScan study and 0.0005 in the UK Biobank Olink study. There was no observed difference in RSA distribution in the surface-exposed region between the full set of missense variants and the 33,445 proteomic GWAS-specific subset [Supplementary Figure 6]. The depletion of buried amino acid residues highlights the proteomic platform focus on soluble proteins, as opposed to transmembrane proteins.

We grouped amino acid residues into three RSA-based classes^20, 21^:

Buried (RSA < 0.2): Residues with minimal solvent exposure, typically located in the protein core.

Surface-exposed (RSA 0.2 – 0.65): Residues with intermediate solvent exposure, which are located on the outer surface of the protein.

Highly accessible (RSA > 0.65): Residues with high solvent accessibility, primarily located in the unstructured regions of the protein.

To test whether platform-discordant missense *cis-*pQTL were more likely to involve a change in positive charge on the protein surface, we performed an enrichment analysis using structural annotations. Specifically, we tested whether substitutions involving the gain or loss of positively charged residues, defined here as arginine, lysine, or histidine, were overrepresented among discordant pQTL located on the structured protein surface (RSA 0.2–0.65). This was then compared against the set of 9,718 proteomic study-identifiable missense SNPs among proteins measured in both proteomics platforms with global MAF > 0.001 and located on surface-exposed regions of their respective proteins. Counts of charge loss and gain were compared between discordant and background sets using a one-sided Fisher’s exact test to determine enrichment.

### ΔΔG

ΔΔG (Delta Delta G) is a measure of the change in free energy difference (ΔG) between the folded and unfolded states of a protein due to a missense mutation in the genome. It is used to assess the impact of missense mutations on protein stability37. ΔΔG values for the missense variants analysed herein were calculated using FoldX version 5.0, an empirical force field-based software designed to predict and analyse the effect of mutations on protein stability and interactions^38^. To compute ΔΔG values, AlphaFold-predicted structures were first pre-processed using the RepairPDB command, after which the BuildModel command was applied to introduce the mutations and estimate their effects on stability.

Although ΔΔG values can be positive (stabilising) or negative (destabilising), they were converted to absolute values to evaluate the extent of conformational changes in the protein’s tertiary structure, regardless of the direction of stability change.

### Data availability

All GWAS summary statistics used in conducting this research are publicly accessible and are available on synapse.org and www.decode.com/summarydata/. The UK Biobank genotypic data used in this study as a LD reference panel were approved under application 19655 and are available to qualified researchers via the UK Biobank data access process. All data compiled and generated in this study regarding proteins and *cis*-pQTL suspected of epitope effects are made available in the supplementary tables. This includes per-platform summaries of protein measurements, harmonized *cis* pQTL lead variants and their high LD associated missense proxies, and their structural annotations.

### Data handling and code availability

Clumping of downloaded summary statistics, variant proxy search and pairwise LD calculation was performed with PLINK (1.9.0p)31.

GWAS results were visualized using the LocusZoom online interface^39^.

Data handling was performed in a Python 3.9.18 environment, with the following primary modules: pandas (2.2.2), scipy (1.11.4), numpy (1.26.4). Plotting was performed using seaborn (0.13.2) and matplotlib (3.9.2). All code used to identify discordant pQTL in this study will be made publicly available upon publication on GitHub (https://github.com/viking-genes/pQTLEpitopes).

## Acknowledgements

J.K. acknowledges the MRC Doctoral Training Programme in Precision Medicine (MR/N013166/1). L.K. was supported by an RCUK Innovation Fellowship from the National Productivity Investment Fund (MR/R026408/1). PN was supported by UKRI’s Medical Research Council (MC_PC_U127592696, MC_PC_U127561128 and MC_UU_00007/10) and the BBSRC (BBS/E/RL/230001A). J.A.M. was supported by funding from the European Research Council (ERC) under the European Union’s Horizon 2020 research and innovation programme (grant agreement No. 101001169). For the purpose of open access, the author has applied a Creative Commons Attribution (CC BY) licence to any Author Accepted Manuscript version arising from this submission.

## Author contributions

Conception and design J.F.W, L.K., J.K.; Data analysis J.K.; Scripting J.K.; Writing J.K.; Data sharing and structural analysis guidance M.B., J.M.; Review and supervision J.F.W., L.K., P.N.

## References

1. Zhao JH, et al. Genetics of circulating inflammatory proteins identifies drivers of immune-mediated disease risk and therapeutic targets. Nature Immunology 24, 1540–1551 (2023).

2. Folkersen L, et al. Genomic and drug target evaluation of 90 cardiovascular proteins in 30,931 individuals. Nature Metabolism 2, 1135–1148 (2020).

3. Ferkingstad E, et al. Large-scale integration of the plasma proteome with genetics and disease. Nat Genet 53, 1712–1721 (2021).

4. Sun BB, et al. Plasma proteomic associations with genetics and health in the UK Biobank. Nature 622, 329–338 (2023).

5. Candia J. SomaScan Bioinformatics: Normalization, Quality Control, and Assessment of Pre-Analytical Variation. bioRxiv, 2024.2002.2009.579724 (2024).

6. Surapaneni A, et al. Identification of 969 protein quantitative trait loci in an African American population with kidney disease attributed to hypertension. Kidney International 102, 1167–1177 (2022).

7. Pozarickij A, et al. Ancestry diversity in the genetic determinants of the human plasma proteome enhances identification of potential drug targets. medRxiv, 2023.2011.2013.23298365 (2025).

8. Xu Y, et al. An atlas of genetic scores to predict multi-omic traits. Nature 616, 123–131 (2023).

9. Haslam DE, et al. Stability and reproducibility of proteomic profiles in epidemiological studies: comparing the Olink and SOMAscan platforms. Proteomics 22, e2100170 (2022).

10. Eldjarn GH, et al. Large-scale plasma proteomics comparisons through genetics and disease associations. Nature 622, 348–358 (2023).

11. Suhre K, et al. Connecting genetic risk to disease end points through the human blood plasma proteome. Nature Communications 8, 14357 (2017).

12. Pietzner M, et al. Mapping the proteo-genomic convergence of human diseases. Science 374, eabj1541 (2021).

13. Joshi A, Mayr M. In Aptamers They Trust. Circulation 138, 2482–2485 (2018).

14. Suhre K, McCarthy MI, Schwenk JM. Genetics meets proteomics: perspectives for large population-based studies. Nature Reviews Genetics 22, 19–37 (2021).

15. Varadi M, et al. AlphaFold Protein Structure Database: massively expanding the structural coverage of protein-sequence space with high-accuracy models. Nucleic Acids Res 50, D439–d444 (2022).

16. Rathore N, et al. Paired Immunoglobulin-like Type 2 Receptor Alpha G78R variant alters ligand binding and confers protection to Alzheimer’s disease. PLOS Genetics 14, e1007427 (2018).

17. Aisina RB, Mukhametova LI. Structure and function of plasminogen/plasmin system. Russian Journal of Bioorganic Chemistry 40, 590–605 (2014).

18. Consortium TU. UniProt: the Universal Protein Knowledgebase in 2025. Nucleic Acids Research 53, D609–D617 (2024).

19. Fischer J, Koblyakova Y, Latendorf T, Wu Z, Meyer-Hoffert U. Cross-Linking of SPINK6 by Transglutaminases Protects from Epidermal Proteases. Journal of Investigative Dermatology 133, 1170–1177 (2013).

20. Wu W, Wang Z, Cong P, Li T. Accurate prediction of protein relative solvent accessibility using a balanced model. BioData Min 10, 1 (2017).

21. Piovesan D, et al. MobiDB: 10 years of intrinsically disordered proteins. Nucleic Acids Res 51, D438–d444 (2023).

22. Davies DR, et al. Unique motifs and hydrophobic interactions shape the binding of modified DNA ligands to protein targets. Proceedings of the National Academy of Sciences 109, 19971–19976 (2012).

23. Potocnakova L, Bhide M, Pulzova LB. An Introduction to B-Cell Epitope Mapping and In Silico Epitope Prediction. Journal of Immunology Research 2016, 6760830 (2016).

24. Rubinstein ND, Mayrose I, Halperin D, Yekutieli D, Gershoni JM, Pupko T. Computational characterization of B-cell epitopes. Molecular Immunology 45, 3477–3489 (2008).

25. Kringelum JV, Nielsen M, Padkjær SB, Lund O. Structural analysis of B-cell epitopes in antibody:protein complexes. Mol Immunol 53, 24–34 (2013).

26. Schweke H, et al. An atlas of protein homo-oligomerization across domains of life. Cell 187, 999-1010.e1015 (2024).

27. Ladner RC. Mapping the epitopes of antibodies. Biotechnology and Genetic Engineering Reviews 24, 1–30 (2007).

28. Hinrichs AS, et al. The UCSC Genome Browser Database: update 2006. Nucleic Acids Research 34, D590–D598 (2006).

29. Karczewski KJ, et al. The mutational constraint spectrum quantified from variation in 141,456 humans. Nature 581, 434–443 (2020).

30. Landini A, et al. Genetic regulation of post-translational modification of two distinct proteins. Nature Communications 13, 1586 (2022).

31. Chang CC, Chow CC, Tellier LC, Vattikuti S, Purcell SM, Lee JJ. Second-generation PLINK: rising to the challenge of larger and richer datasets. GigaScience 4, (2015).

32. Miller S, Janin J, Lesk AM, Chothia C. Interior and surface of monomeric proteins. J Mol Biol 196, 641–656 (1987).

33. Agirre J, et al. The CCP4 suite: integrative software for macromolecular crystallography. Acta Crystallogr D Struct Biol 79, 449–461 (2023).

34. Tunyasuvunakool K, et al. Highly accurate protein structure prediction for the human proteome. Nature 596, 590–596 (2021).

35. Auton A, et al. A global reference for human genetic variation. Nature 526, 68–74 (2015).

36. Sherry ST, et al. dbSNP: the NCBI database of genetic variation. Nucleic Acids Research 29, 308–311 (2001).

37. Fersht AR, Matouschek A, Serrano L. The folding of an enzyme. I. Theory of protein engineering analysis of stability and pathway of protein folding. J Mol Biol 224, 771–782 (1992).

38. Delgado J, Radusky LG, Cianferoni D, Serrano L. FoldX 5.0: working with RNA, small molecules and a new graphical interface. Bioinformatics 35, 4168–4169 (2019).

39. Boughton AP, et al. LocusZoom.js: interactive and embeddable visualization of genetic association study results. Bioinformatics 37, 3017–3018 (2021).

